# Quantifying the Information Capacity of DNA Methylation as an Epigenetic Memory System

**DOI:** 10.64898/2026.06.28.735086

**Authors:** Ildefonso M. De la Fuente, José Carrasco-Pujante, María Fedetz, Leire Legarreta, Iker Malaina, Borja Camino-Pontes, Gorka Pérez-Yarza, Luis Martínez, Jesús M Cortés, José I. López

**Affiliations:** Department of Nutrition, CEBAS-CSIC Institute, Espinardo University Campus, 30100 Murcia, Spain; Department of Mathematics, Faculty of Science and Technology, University of the Basque Country, UPV/EHU, 48940 Leioa, Spain; Department of Cell Biology and Histology, Faculty of Medicine and Nursing, University of the Basque Country, UPV/EHU, 48940 Leioa, Spain; Department of Cell Biology and Immunology, Institute of Parasitology and Biomedicine “López-Neyra”, CSIC, 18016 Granada, Spain; Biobizkaia Health Research Institute, 48903 Barakaldo, Spain; IKERBASQUE: The Basque Foundation for Science, 48009 Bilbao, Spain

**Keywords:** DNA methylation, epigenetic processes, methylome information capacity, CpG methylation, Shannon entropy

## Abstract

The information content of the genome has been extensively analyzed. However, a comparable quantitative framework for DNA methylation is still lacking. Without such quantification, the magnitude of this regulatory and dynamic epigenetic structure remains conceptually imprecise, even though methylation dysregulation is strongly linked to disease-related phenotypes and altered cellular identity. Here we address this gap by applying Shannon information theory to DNA methylation. We first consider methylation marks as binary or probabilistic regulatory states and estimate the theoretical upper-bound information capacity of the human methylome under simplifying assumptions. We then progressively refine this estimate by incorporating biologically relevant constraints, including methylation bias, bimodal methylation distributions, local CpG correlation, genomic regulatory class, and cell-type-discriminative methylation patterns. This approach allows us to distinguish between theoretical methylation capacity, statistical methylation entropy, and biologically interpretable regulatory information. Finally, we consider methylation information from a discriminative perspective, analyzing its contribution to distinguishing cell types and regulatory cellular states. Within this framework, mutual information between methylation patterns and cell identity provides a biologically constrained estimate of methylation’s role as an epigenetic identity code. Our layered analysis reconciles megabit-scale methylome capacity with compact, biologically interpretable identity signatures.

**Graphical Abstract:** 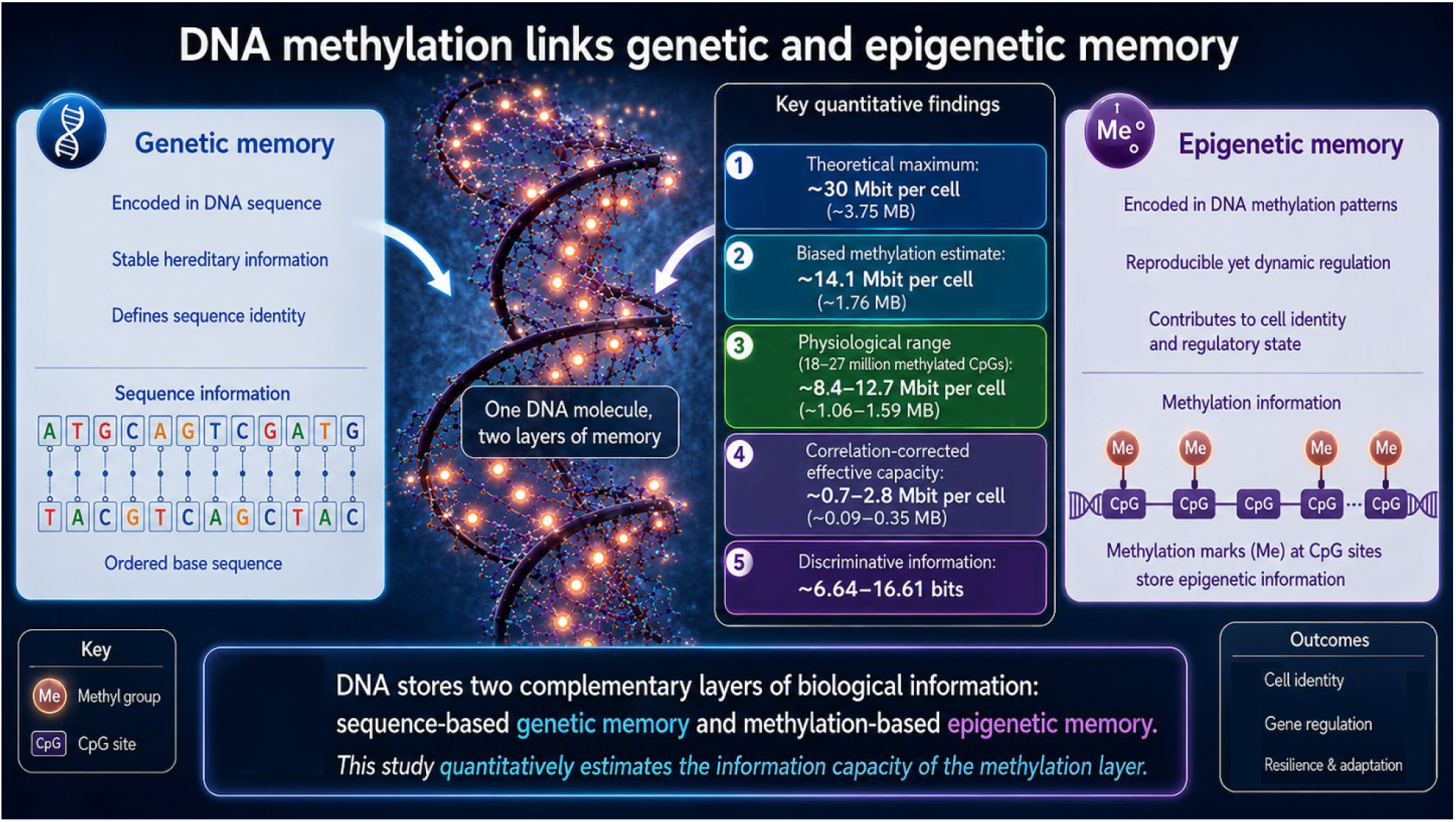

## 1. Introduction

DNA methylation is one of the most fundamental and widely studied epigenetic modifications (1–3). By adding a methyl group to cytosine residues, most commonly at CpG dinucleotides in mammals, cells acquire a covalent regulatory layer superimposed on the primary DNA sequence. Unlike genetic information, DNA methylation does not alter the identity of nucleotides. Instead, it modulates chromatin organization, transcriptional competence, genome stability, repression of transposable elements, imprinting, developmental programs, and cell identity. Because methylation patterns can be propagated through cell division and, at the same time, be reversible during development, reprogramming, aging, and disease, DNA methylation is often considered a molecular substrate of epigenetic memory.

Several characteristics make DNA methylation particularly relevant as an information-carrying regulatory system. First, methylation marks are chemically stable but are dynamically regulated by the opposing activities of methyltransferases and demethylation pathways (4, 5). Second, methylation patterns are highly structured across the genome rather than randomly distributed, with distinct behaviors at promoter CpG islands, CpG island shores, enhancers, gene bodies, imprinting control regions, repetitive elements, and transposable elements (2, 6, 7). Third, methylation is highly dependent on cell type and lineage, allowing cells with the same genome to maintain different regulatory identities. Finally, aberrant methylation patterns are widely associated with pathological phenotypes, including developmental disorders, aging-related changes, cancer, metabolic disease, and other disease-associated transcriptional states. These properties suggest that DNA methylation is not merely a biochemical modification, but a dynamic regulatory architecture capable of storing, transmitting, and constraining biological information.

Despite significant progress in cataloging DNA methylation landscapes across tissues, developmental stages, cell types, and pathological states (7–10), a central quantitative question remains insufficiently addressed: what is the potential information capacity of these biochemical covalent marks? The information content of the genome has been extensively discussed in relation to nucleotide sequence, coding potential, regulatory annotation, and large-scale functional genomics efforts such as ENCODE (11). However, a comparable quantitative framework for DNA methylation is still lacking. Without such quantification, the magnitude of this regulatory and dynamic epigenetic structure remains conceptually imprecise, even though methylation dysregulation is strongly linked to disease-related phenotypes and altered cellular identity.

Here we address this gap by applying Shannon information theory to DNA methylation (12, 13). We consider methylation marks as binary or probabilistic regulatory states and estimate the theoretical upper-bound information capacity of the human methylome under simplifying assumptions. We then progressively refine this estimate by incorporating biologically relevant constraints, including methylation bias, bimodal methylation distributions, local CpG correlation, genomic regulatory class, and cell-type-discriminative methylation patterns. This approach allows us to distinguish between theoretical methylation capacity, statistical methylation entropy, and biologically interpretable regulatory information.

Using approximately 30 million CpG sites as an upper-bound reference in human cells, we estimate a theoretical maximum capacity of approximately 30 Mbit per cell under an idealized binary model in which each CpG behaves as an independent methylated/unmethylated variable with maximal entropy. Under a more conservative methylation-biased model, in which the average entropy per CpG is reduced, the estimated capacity falls to the megabit range. Considering a physiological range corresponding to approximately 18–27 million methylated CpG-equivalent states, the entropy-corrected capacity is estimated at approximately 8.44–12.66 Mbit per cell. These values should not be interpreted as the amount of biologically usable information in the methylome, but as upper-bound or entropy-corrected estimates of the potential space of regulatory states.

Furthermore, we show that the effective regulatory information of DNA methylation is substantially compressed by genomic organization and biological interpretation. CpG sites are not independent informational units; neighboring CpGs frequently behave as locally correlated methylation blocks. Under an idealized independent-site model, 30 million CpGs with a conservative entropy of ~0.47 bits per CpG yield approximately 14.07 Mbit of methylation information. However, when local CpG correlation is introduced, this value decreases markedly: correlation lengths of 5, 10, and 20 CpGs per effective methylation unit reduce the estimate to approximately 2.81 Mbit, 1.41 Mbit, and 0.70 Mbit, respectively. Thus, local methylation coupling compresses the apparent genome-wide information capacity by roughly one to two orders of magnitude, producing a lower but more biologically plausible range of approximately 0.70–2.81 Mbit per cell. We also show that methylation information is not uniformly distributed across the genome. Enhancers, CpG island shores, imprinting control regions, lineage-defining differentially methylated regions, and other regulatory elements may carry disproportionate biological information despite representing only a subset of methylated cytosines.

Finally, we consider methylation information from a discriminative perspective, analyzing its contribution to distinguishing cell types and regulatory cellular states. Within this framework, mutual information between methylation patterns and cell identity provides a biologically constrained estimate of methylation’s role as an epigenetic identity code. This discriminative information is conditioned by the entropy of the cell state space and therefore depends on the resolution at which cellular identity is defined. Thus, methylation can simultaneously possess megabit-scale theoretical capacity and much smaller, highly compressed, biologically meaningful discriminative information.

Taken together, this work provides a quantitative framework for understanding DNA methylation as an epigenetic memory system. By estimating its theoretical, statistical, correlation-corrected, and cell-type-discriminative information content, we place DNA methylation within an information-theoretic framework that can be refined using methylome atlases, single-cell methylation data, disease methylomes, and regulatory annotations. Such quantification may help clarify the magnitude of methylation-based regulation, improve comparisons between normal and pathological epigenetic states, and support future quantitative studies of methylation dysregulation in disease-related phenotypes (14–16). All numerical results shown here are fully reproducible using the “Python Reproducibility Modules” provided in Supplementary Information S4.

## 2. Methods

### 2.1. Conceptual scope

This study develops an information-theoretic framework to estimate the potential statistical information capacity of DNA methylation in the human methylome. The approach is grounded in Shannon information theory, where information is interpreted as reduction of uncertainty, and in the distinction between statistical entropy and biologically decoded regulatory meaning (12, 13). DNA methylation is treated as a stable yet reversible epigenetic modification that contributes to cellular identity, genome regulation, and epigenetic memory (1–4).

The analysis defines a hierarchy of increasingly constrained estimates: a binary upper bound, an entropy-corrected estimate that incorporates methylation bias, a methylation-burden-adjusted approximation, a correlation-corrected estimate based on effective methylation units, a regulatory-class-stratified formulation, and a discriminative information framework for cell-state resolution. This structure reflects the known context-dependence of DNA methylation across CpG islands, CpG island shores, enhancers, gene bodies, imprinting control regions, repetitive elements, and transposable elements (2, 6, 7, 11).

Throughout the framework, numerical values are interpreted as order-of-magnitude estimates under explicit assumptions. This is essential because CpG methylation is biased, locally correlated, developmentally constrained, cell-type dependent, and interpreted through broader chromatin and regulatory contexts (8, 9, 17, 18).

### 2.2. CpG reference space and theoretical upper-bound capacity

The human methylome was approximated using an upper-bound reference of ~30 million CpG sites. In the idealized binary model, each CpG site can exist in one of two methylation states: methylated or unmethylated. For N CpG sites, the total number of possible methylation configurations is therefore:

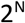

Under maximal uncertainty, where methylated and unmethylated states are equally probable, each CpG contributes 1 bit of information (12, 13).

Using N_CpG_ = 30 × 10^6^ CpG sites, the theoretical upper-bound methylation capacity was calculated as:

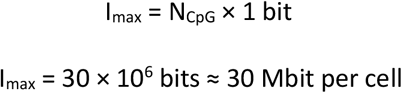

This estimate assumes complete independence among CpG sites and maximal entropy at every site. It is therefore not interpreted as biologically usable methylation information, but rather as the maximal binary state space under idealized assumptions.

### 2.3. Shannon entropy of a biased CpG methylation variable

To account for methylation bias, CpG methylation was modelled as a Bernoulli random variable with methylation probability p. The Shannon entropy per CpG site was calculated as (12, 13):

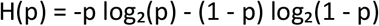

Here, H(p) is the entropy per CpG site, p is the probability of methylation, and 1 - p is the probability of being unmethylated. Entropy is maximal when p = 0.5, giving H(p) = 1 bit per CpG. Mammalian methylation profiles are generally not balanced around p = 0.5; many CpGs are either highly methylated or largely unmethylated, producing biased and often bimodal methylation distributions (2, 8, 9).

A conservative biased methylation scenario was evaluated using p = 0.9. Under this assumption:

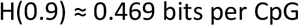

For readability, this value was rounded to approximately 0.47 bits per CpG in the manuscript text and figures. The entropy-corrected methylome capacity was calculated as:

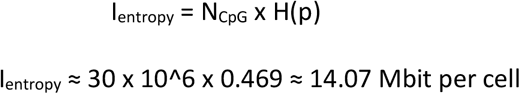

This estimate accounts for methylation bias but still assumes that CpG sites behave as statistically independent methylation variables.

### 2.4. Methylation-burden-adjusted capacity estimate

To provide a physiologically constrained approximation, the calculation was also restricted to a methylation-burden range equivalent to approximately 18-27 million methylatable CpG sites. This range was treated as a burden-adjusted approximation of the methylated component of the methylome, not as a direct count of independent informative CpG variables.

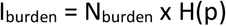

where N_burden_ represents the physiological methylation-burden range and H(p) is the entropy per CpG under the biased methylation model. Using N_burden_ = 18-27 x 10^6 and H(p) ≈ 0.469 bits:

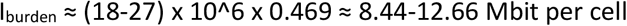

This calculation should be interpreted as a methylation-burden-adjusted approximation rather than a direct count of independent informative CpG variables.

### 2.5. Correlation-corrected effective methylation units

The independent-site assumption is biologically unrealistic because neighboring CpGs often show correlated methylation states. Local methylation coupling and CpG topology influence methylation patterns across genomic regions, reducing the effective number of independent methylation variables (17). DNA-binding factors and distal regulatory context also shape local methylation landscapes (18).

To account for local CpG coupling, the methylome was represented as a set of effective methylation units, each containing an average number of locally correlated CpGs. This average local correlation length was denoted as L_c_.

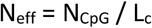

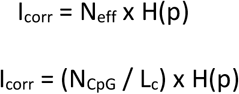

where N_CpG_ is the total CpG reference space, L_c_ is the assumed average local correlation length, N_eff_ is the effective number of independent methylation units, and H(p) is the entropy per unit. A sensitivity analysis was performed using L_c_ values of 1, 5, 10, and 20 CpGs per effective methylation unit.

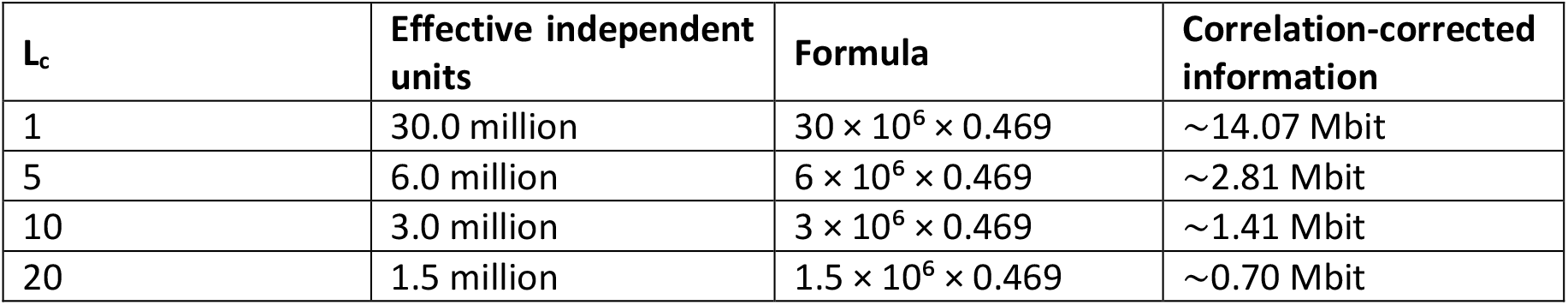

| $L_c$ | Effective independent units | Formula | Correlation-corrected information |
| --- | --- | --- | --- |
| 1 | 30.0 million | $30 \times 10^6 \times 0.469$ | ~14.07 Mbit |
| 5 | 6.0 million | $6 \times 10^6 \times 0.469$ | ~2.81 Mbit |
| 10 | 3.0 million | $3 \times 10^6 \times 0.469$ | ~1.41 Mbit |
| 20 | 1.5 million | $1.5 \times 10^6 \times 0.469$ | ~0.70 Mbit |

This analysis illustrates how local CpG correlation compresses the effective methylation state space by reducing the number of statistically independent methylation units. The selected L_c values are not assumed to be universal constants; they are used as an order-of-magnitude sensitivity analysis that can be replaced by empirical correlation lengths in future methylome datasets.

### 2.6. Regulatory-class-stratified methylation entropy

A genome-wide estimate assumes that all CpGs contribute equally to methylation information, but this assumption is not biologically justified. CpGs located in promoter CpG islands, CpG island shores, enhancers, imprinting control regions, repeats, transposable elements, and gene bodies differ in methylation probability, variability, cell-type specificity, and regulatory interpretation (2, 6, 7, 11, 19).

For each genomic regulatory class g, the information contribution was expressed as:

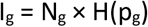

where N_g_ is the number of CpGs or effective methylation units assigned to class g, p_g_ is the methylation probability within that class, and H(p_g_) is the corresponding Shannon entropy. The total statistical methylation information across regulatory classes can then be expressed as:

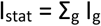

This formulation separates genome-wide methylation variability from regulatory interpretation. It also emphasizes that the biological value of methylation information depends not only on the number of CpGs involved, but also on genomic location, chromatin state, regulatory context, cell-type specificity, and functional interpretability (7, 9).

### 2.7. Discriminative methylation information and cellular-state resolution

In addition to estimating methylation information as a property of the methylome, the framework considers how methylation patterns may discriminate among cell types or regulatory cellular states. This was formulated using mutual information between methylation pattern M and cellular identity C. This formulation follows the Shannon information-theoretic interpretation of information as uncertainty reduction (12, 13).

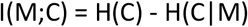

where H(C) is the uncertainty about cellular identity before observing methylation, and H(C|M) is the remaining uncertainty after methylation information is known. Because mutual information is bounded by the entropy of the decoded variable:

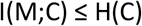

the maximum discriminative information depends on how the cellular-state space is defined. If cellular identity is represented as K equally probable states, then:

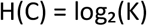

This distinction is important because methylation atlases and single-cell epigenomic studies show that methylation patterns can discriminate cellular identity at multiple biological resolutions, from broad cell classes to finer regulatory states (10, 20, 21).

### 2.8. Hierarchical classification of methylation-defined cellular states

The discriminative variable C can be defined at multiple levels of biological resolution. In the simplest case, C may represent a flat cell-type label. However, this representation is biologically coarse because cellular identity is not only determined by broad cell type, but also by tissue origin, developmental lineage, subtype, activation state, differentiation state, disease-associated state, epigenetic age, and allele-specific regulatory status. Current cell taxonomy and ontology frameworks explicitly emphasize that cell identity includes both cell types and cell states, motivating a hierarchical or combinatorial representation of the decoded biological variable (22–24).

To represent this hierarchy, the cellular identity variable was generalized as:

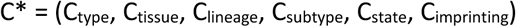

where C* denotes a methylation-defined regulatory cellular state rather than a single flat cell-type label. Under this formulation, broad cell-type identity is only one component of the decoded state space. Additional components capture tissue context, lineage history, cellular subtype, functional or activation state, developmental stage, disease-associated state, and imprinting or allele-specific regulatory status.

The entropy of this hierarchical cellular identity variable can be decomposed using conditional entropy terms:

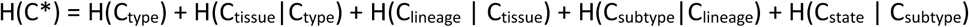

This decomposition does not require each component to be independent. Instead, it recognizes that the uncertainty contributed by each biological layer is conditional on the preceding layers. For example, the uncertainty about tissue context may depend on the broad cell type, and the uncertainty about subtype may depend on lineage and tissue. This conditional formulation is therefore more biologically plausible than treating all identity descriptors as independent categorical variables.

Within this framework, the discriminative information estimate becomes:

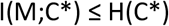

rather than being restricted to:

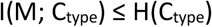

This distinction is central to the interpretation of methylation information. A low-bit estimate for a coarse classification problem, such as distinguishing 100 broad cell types, does not imply that the methylome contains little information. It means only that few reliable bits are sufficient to identify one label among a limited set of alternatives. At higher biological resolution, methylation patterns may encode or preserve information about tissue-specific enhancer states, lineage memory, developmental transitions, imprinting status, epigenetic age, disease-associated remodeling, and regulatory-state diversity (7, 10, 20, 21).

The hierarchical formulation therefore provides a bridge between genome-wide methylation entropy and cell-state-discriminative methylation information. Genome-wide methylation entropy can reach the megabit scale because it refers to possible methylation states across millions of CpGs or effective methylation units. By contrast, discriminative information is bounded by the entropy of the biological variable being decoded. These two quantities are not contradictory because they measure different random variables.

### 2.9. Sensitivity analysis of discriminative information

A sensitivity analysis was performed to illustrate how upper-bound discriminative information changes as the number of distinguishable methylation-defined cellular states increases. For K equally probable states, the maximal uncertainty of the cellular-state variable was calculated as:

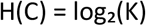

Four illustrative values of K were evaluated: 100, 1,000, 10,000, and 100,000 states. These values correspond to increasingly resolved biological classification spaces, from a broad cell-type atlas to a combinatorial space incorporating cell type, tissue context, developmental stage, functional state, or disease-associated regulatory state.

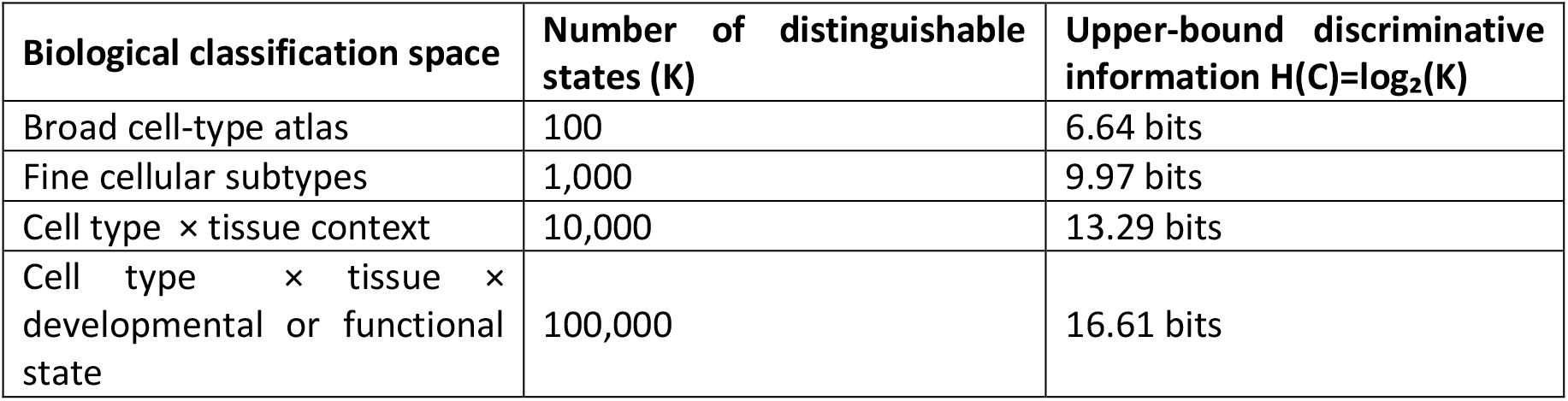

This sensitivity analysis is not intended to assert that all of these states have already been experimentally resolved by methylation alone. Rather, it shows how the upper bound of I(M;C) changes when the decoded biological state space is represented at different resolutions. Thus, the 6.64-bit value for 100 cell types should be interpreted as a lower-resolution discriminative benchmark, not as the total information capacity of DNA methylation.

The analysis also clarifies why methylation can simultaneously display megabit-scale genome-wide entropy and compact bit-scale discriminative information. The former quantifies a potential methylation state space distributed across many genomic sites, whereas the latter quantifies uncertainty reduction about a defined biological label or state variable.

### 2.10. Interpretation of model layers and limitations

The estimates generated by this framework correspond to different layers of interpretation. The theoretical maximum describes an idealized binary state space. The entropy-corrected estimate accounts for biased methylation probabilities. The methylation-burden-adjusted estimate restricts the calculation to a physiologically plausible burden range. The correlation-corrected estimate accounts for local CpG dependency. The regulatory-class-stratified formulation introduces genomic context. The discriminative information estimate asks how much methylation can reduce uncertainty about a defined cellular-state variable.

Several limitations follow from these assumptions. First, the number of CpGs, methylation burden, methylation probability, and local correlation length vary across tissues, developmental stages, organisms, cell types, and assay platforms. Second, a binary methylated/unmethylated model simplifies biochemical complexity, including hydroxymethylcytosine, oxidized cytosine derivatives, allele-specific methylation, strand asymmetry, non-CpG methylation, and single-cell heterogeneity. Third, methylation information is interpreted by chromatin state, transcription factor binding, replication timing, three-dimensional genome organization, and broader gene-regulatory networks. Therefore, methylation information should not be treated as an autonomous code independent of the epigenomic system (2, 7, 9, 19).

### 2.11. Reproducibility of numerical estimates

All numerical estimates were generated from the equations described above using reproducible Python modules provided in the Supplementary Information. These modules calculate Shannon entropy for different methylation probabilities, upper-bound methylome capacity, methylation-burden-adjusted estimates, correlation-corrected effective methylation units, and discriminative information as a function of cellular-state resolution.

The reproducibility modules are intended to allow modification of the main parameters, including the number of CpG sites, methylation probability, physiological methylation-burden range, local correlation length, and number of distinguishable cellular states. This allows the framework to be adapted to future empirical methylome datasets, single-cell methylation profiles, disease methylomes, or organism-specific methylation landscapes.

## 3. Results

In this work, information capacity is used in a statistical and information-theoretic sense, not as a direct measure of biologically decoded regulatory meaning. The theoretical capacity estimate assumes that CpG methylation can be represented as a binary variable and that CpG sites are independent. Subsequent estimates progressively relax this assumption by incorporating methylation bias, local CpG correlation, regulatory genomic class, and cell-state discrimination. Therefore, the numerical values presented here should be interpreted as order-of-magnitude estimates under explicitly defined assumptions rather than fixed biological constants.

### 3.1. Theoretical and entropy-corrected methylome capacity

DNA methylation defines a constrained, heritable regulatory state space whose theoretical information capacity can be estimated using Shannon entropy. In a simplified binary model, each CpG site can exist in one of two states, methylated or unmethylated. For N CpG sites, the number of possible methylation configurations is therefore 2^N^. Taking approximately 30 million CpG sites as an upper-bound reference for the human methylome, this corresponds to a theoretical maximum capacity of approximately 30 Mbit per cell, assuming that each CpG behaves as an independent binary variable with maximal entropy.

For a CpG site with methylation probability p, the Shannon entropy (12, 13) is:

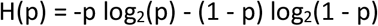

The maximum value, H(p) = 1 bit, occurs when p = 0.5. However, mammalian CpG methylation is not generally balanced around 50%. Many CpGs are either highly methylated or largely unmethylated, producing biased and often bimodal methylation distributions (2, 8, 9). Under a conservative methylation-biased model with p = 0.9, the entropy per CpG is:

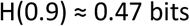

Using this entropy value, 30 million CpGs yield an entropy-corrected estimate of approximately:

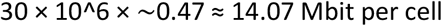

If the calculation is restricted to a physiological methylation-burden range equivalent to approximately 18–27 million methylatable CpG sites, the estimated capacity becomes:

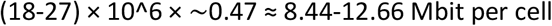

This calculation should be interpreted as a methylation-burden-adjusted approximation rather than a direct count of independent informative CpG variables (Table 1).

**Table 1.**
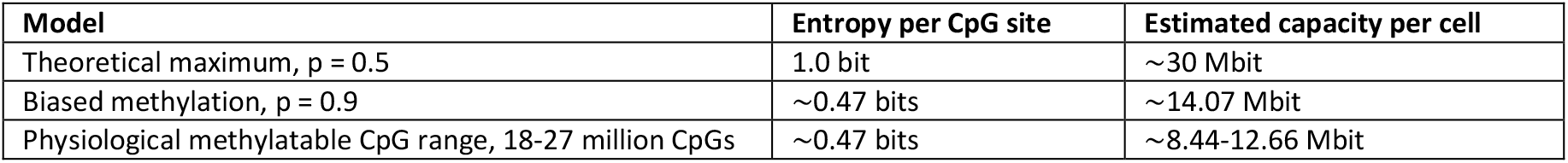
Information capacity of DNA methylation.

| Model | Entropy per CpG site | Estimated capacity per cell |
| --- | --- | --- |
| Theoretical maximum, $p = 0.5$ | 1.0 bit | ~30 Mbit |
| Biased methylation, $p = 0.9$ | ~0.47 bits | ~14.07 Mbit |
| Physiological methylatable CpG range, 18-27 million CpGs | ~0.47 bits | ~8.44-12.66 Mbit |

These values define the first level of the framework: the methylome has megabit-scale theoretical and entropy-corrected capacity, although this capacity is expected to be substantially reduced by biological constraints (Figure 1).

**Figure 1.**
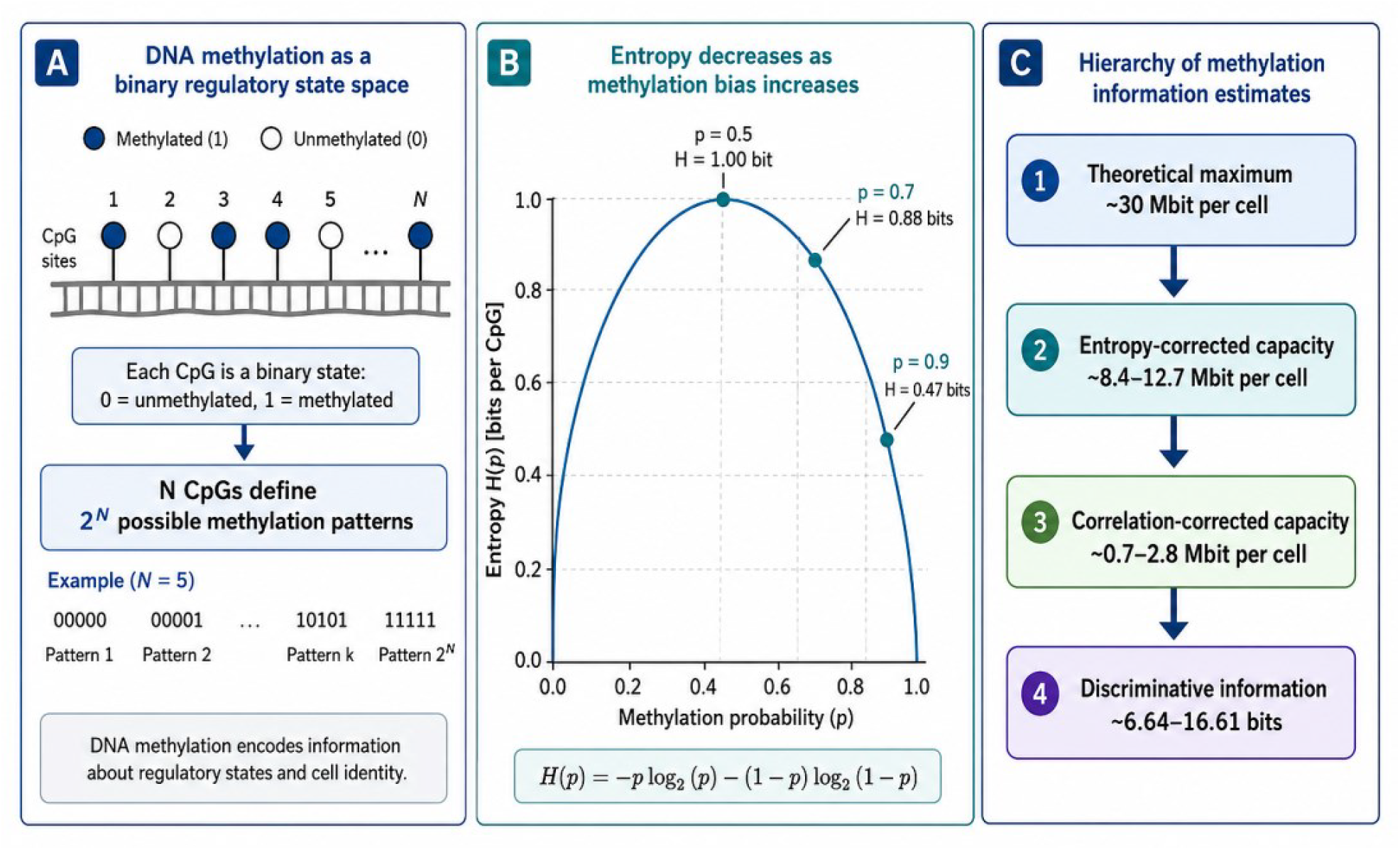
Information-theoretic framework of DNA methylation as an epigenetic memory system. (A) Conceptual representation of CpG methylation as a binary regulatory state space, in which each CpG site can be considered methylated (1) or unmethylated (0). Under an idealized independent-site model, *N*_CpGs define 2^N possible methylation configurations. (B) Shannon entropy of a binary methylation variable as a function of methylation probability, *p*. Entropy is maximal at *p* = 0.5, where H(*p*) = 1 bit per CpG, and decreases as methylation becomes biased toward either the methylated or unmethylated state. Representative values are shown for *p* = 0.7 and *p* = 0.9. (C) Hierarchical interpretation of methylation information estimates. The theoretical maximum capacity of the human methylome is approximately 30 Mbit per cell, whereas methylation bias, local CpG correlation, and biological interpretation progressively reduce the effective information content. This framework distinguishes theoretical capacity, entropy-corrected capacity, correlation-corrected capacity, and discriminative information related to cellular identity.

### 3.2. Effective regulatory information and biological compression

The theoretical estimates expressed above treat CpG sites as independent informational units. In living cells, however, this assumption is biologically unrealistic. CpG methylation is biased, locally correlated, developmentally constrained, and interpreted through genomic regulatory context (9, 17, 18). Therefore, the biologically effective information content of DNA methylation is expected to be lower than the binary or entropy-corrected upper bounds.

Three levels of methylation information can be distinguished. First, theoretical information refers to the maximal binary capacity obtained when CpGs are treated as independent methylated/unmethylated variables. Second, statistical information refers to the entropy observable after accounting for methylation bias, bimodality, and local correlation. Third, functional regulatory information refers to the subset of methylation variation that is reproducible, cell-type-specific, associated with regulatory genomic elements, and interpretable in relation to transcriptional, chromatin, developmental, or allele-specific outcomes.

Accordingly, DNA methylation should not be interpreted as an unconstrained digital storage system. Rather, it behaves as a structured and compressed regulatory memory. Most CpGs are constitutively methylated or unmethylated within a given cellular context, whereas a smaller subset of regulatory regions, including enhancers, CpG island shores, imprinting control regions, and lineage-specific differentially methylated regions, carries disproportionate biological information.

### 3.3. Entropy stratified by genomic regulatory class

A genome-wide estimate assumes that all CpGs contribute equally to methylation information. This assumption is not biologically justified. CpGs located in promoter CpG islands, CpG island shores, enhancers, imprinting control regions, repetitive elements, and gene bodies differ in methylation probability, variability, cell-type specificity, and regulatory interpretation (2, 6, 7, 11).

For each genomic class g, the information contribution can be approximated as:

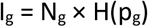

where N_g_ is the number of CpGs or effective methylation units assigned to class g, p_g_ is the methylation probability within that class, and H(p_g_) is the corresponding entropy. The total statistical methylation information can therefore be expressed as:

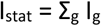

This formulation separates raw methylation variability from biologically interpretable regulatory information (Table 2).

**Table 2.** Conceptual regulatory-class decomposition of methylation information.

| Genomic class | Typical methylation behavior | Expected entropy | Functional interpretation |
| --- | --- | --- | --- |
| Promoter CpG islands | Often constitutively unmethylated in normal cells | Low under physiological conditions | Low variability, but high functional consequence when developmentally or aberrantly methylated |
| CpG island shores | More variable than CpG island cores | Moderate to high | Informative for tissue specificity and developmental transitions |
| Enhancers | Frequently cell-type-specific and dynamically methylated | High in regulatory subsets | Important for cell identity and transcription factor accessibility |
| Imprinting control regions | Parent-of-origin-specific and mitotically stable | Low genome-wide abundance, high locus-specific information | Encodes allele-specific developmental regulation |
| Repeats and transposable elements | Usually highly methylated and repressed | Low to moderate | Genome defense and repression |
| Gene bodies | Often methylated in active genes, context-dependent | Moderate | May contribute to transcriptional stability and suppression of cryptic initiation |

This stratified framework emphasizes that the functional value of methylation information depends not only on the number of CpGs, but also on their genomic location and regulatory meaning.

### 3.4. Correlation-corrected effective methylation units

CpG methylation states are locally correlated. Neighboring CpGs within CpG islands, shores, enhancers, imprinting control regions, repeats, and partially methylated domains often behave as coordinated blocks. As a result, the number of statistically independent methylation variables is smaller than the total number of CpG sites.

A simple correlation-corrected estimate can be written as:

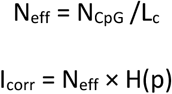

or equivalently:

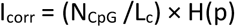

where N_CpG_ is the total number of CpG sites considered, L_c_ is the average local correlation length expressed as CpGs per effective methylation unit, N_eff_ is the effective number of independent methylation units, and H(p) is the entropy per unit.

Using N_CpG_ = 30 million and H(p) = ~0.47 bits, the sensitivity of the estimate to local correlation is shown below.

Thus, if CpGs are treated as independent, the biased entropy estimate is approximately 14.07 Mbit. However, if methylation is locally correlated in blocks of 5-20 CpGs, the estimate decreases to approximately 0.70-2.81 Mbit per cell (Table 3, Figure 2). This range should not be interpreted as a universal constant, but as an order-of-magnitude sensitivity analysis showing how local CpG coupling compresses the effective regulatory state space.

**Table 3.** Correlation-corrected methylation information.

| Average local correlation length, $L_c$ | Effective independent units, $N_{\text{eff}}$ | Correlation-corrected information |
| --- | --- | --- |
| 1 CpG per unit | 30.0 million | $\sim 14.07$ Mbit |
| 5 CpGs per unit | 6.0 million | $\sim 2.81$ Mbit |
| 10 CpGs per unit | 3.0 million | $\sim 1.41$ Mbit |
| 20 CpGs per unit | 1.5 million | $\sim 0.70$ Mbit |

**Figure 2.**
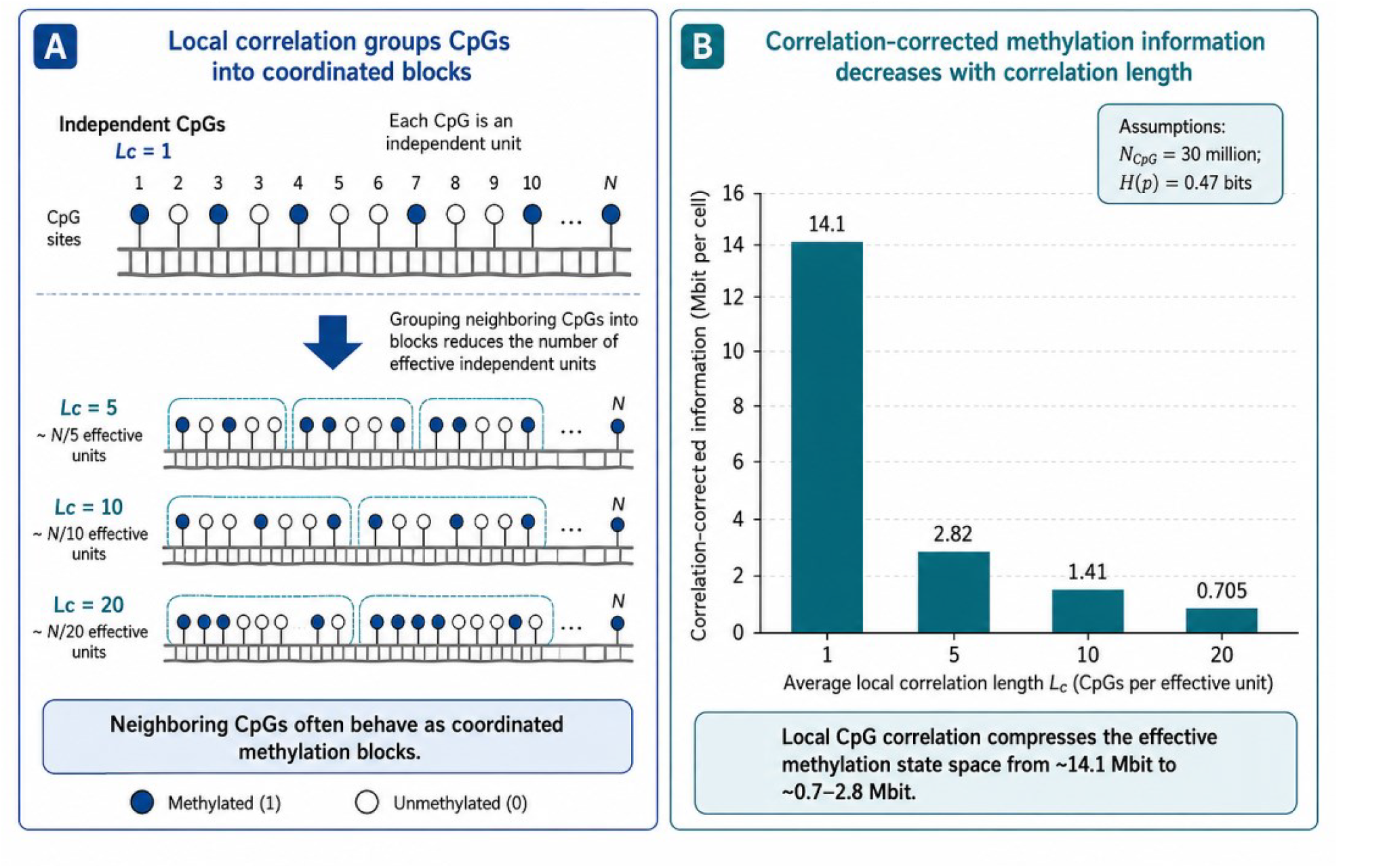
Compression of methylation information by local CpG correlation. (A) Schematic illustration of local CpG correlation. In the independent-site model, each CpG is treated as a separate informational unit. In biological methylomes, neighboring CpGs often behave as coordinated methylation blocks, reducing the effective number of independent methylation units. Increasing the average local correlation length, L_c_, therefore compresses the methylation state space from N_CpGs_ to approximately N/L_c_ effective units. (B) Sensitivity analysis of correlation-corrected methylation information using *N*CpG = 30 million and H(*p*) = ~0.47 bits per CpG. Under an independent-site assumption (L_c_ = 1), the biased entropy estimate is approximately 14.07 Mbit per cell. When local correlation lengths of 5, 10, and 20 CpGs per effective unit are assumed, the estimate decreases to approximately 2.81, 1.41, and 0.70 Mbit per cell, respectively. These values illustrate how local methylation coupling compresses the effective regulatory information capacity of the methylome.

### 3.5. Discriminative methylation information between cell types

The preceding estimates quantify methylation information as a property of the methylome itself. A more biologically constrained question is how much methylation contributes to distinguishing cell types or regulatory cellular states. This can be formulated using the mutual information between methylation pattern M and cellular identity C (10, 12, 13):

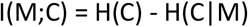

where H(C) is the uncertainty about cellular identity before observing methylation, and H(C|M) is the remaining uncertainty after methylation information is known. Equivalently, mutual information measures how much knowing the methylation pattern reduces uncertainty about the cellular state.

This quantity is bounded by the entropy of the cellular identity variable:

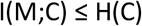

If cellular identity is represented as K equally probable states, then:

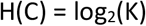

These values are much smaller than the megabit-scale genome-wide capacity estimates because they answer a different question (Table 4). Genome-wide entropy estimates quantify the potential state space of methylation patterns distributed across millions of CpGs or effective methylation units. By contrast, discriminative information quantifies the number of bits required to identify one biological state among a defined set of alternatives (Figure 3).

**Table 4.** Upper-bound discriminative information as a function of cellular-state resolution.

| Biological classification space | Number of distinguishable states, $K$ | Upper-bound discriminative information |
| --- | --- | --- |
| Broad cell-type atlas | 100 | 6.64 bits |
| Fine cellular subtypes | 1,000 | 9.97 bits |
| Cell type × tissue context | 10,000 | 13.29 bits |
| Cell type × tissue × developmental or functional state | 100,000 | 16.61 bits |

**Figure 3.**
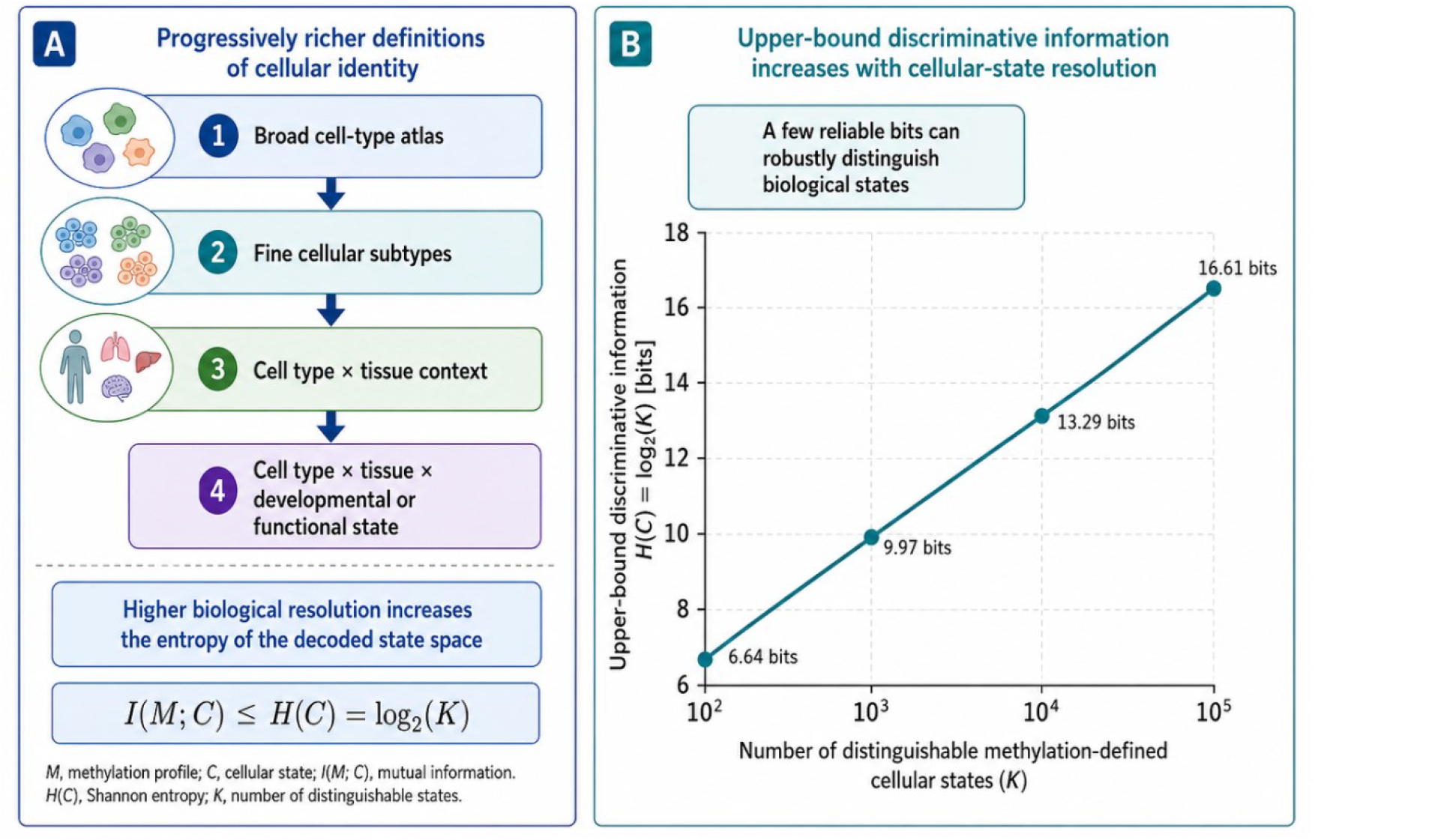
Discriminative methylation information depends on cellular-state resolution. (A) Conceptual hierarchy of cellular identity definitions. Methylation patterns may discriminate broad cell-type labels, finer cellular subtypes, tissue-specific contexts, or more complex regulatory states incorporating developmental, functional, or disease-associated features. The discriminative information carried by methylation is bounded by the entropy of the cellular-state variable, such that I(*M*;*C*) ≤ H(*C*) = log_2_(*K*), where *M* is the methylation profile, *C* is cellular identity, and *K* is the number of distinguishable methylation-defined states. (B) Upper-bound discriminative information as a function of cellular-state resolution. For 100, 1,000, 10,000, and 100,000 equally probable states, H(*C*) corresponds to 6.64, 9.97, 13.29, and 16.61 bits, respectively. These bit-scale values should not be interpreted as the total information capacity of the methylome, but as the information required to distinguish one biological state among a defined set of alternatives.

Therefore, the 6.64-bit value for 100 cell types should not be interpreted as the total information stored in the methylome. These estimates are not contradictory because they measure different random variables: genome-wide methylation entropy quantifies the potential methylation state space, whereas discriminative information quantifies uncertainty reduction about a defined cellular-state variable. It is the upper bound for a coarse classification task. At higher biological resolution, including tissue origin, lineage, subtype, activation state, developmental stage, disease-associated state, or imprinting status, the entropy of the cellular identity variable increases.

This distinction reconciles two apparently different scales of methylation information. DNA methylation can exhibit megabit-scale theoretical and entropy-corrected capacity while also functioning as a compact, highly reliable identity code. The biologically most relevant methylation information is therefore not distributed uniformly across all CpGs, but concentrated in reproducible, cell-type-specific, and regulatory methylation features that help define cellular identity and epigenetic memory.

## 4. Discussion

DNA methylation is widely recognized as a central component of epigenetic regulation, cellular identity, genome stability, and heritable transcriptional control (1, 2, 4). However, despite extensive progress in mapping methylation landscapes across tissues, cell types, developmental stages, and pathological states, the magnitude of methylation as an information-bearing system has remained conceptually ill-defined. In this work, we addressed this gap by applying Shannon information theory to DNA methylation and by progressively constraining the estimate from an idealized binary methylation model toward more biologically realistic regulatory representations (9, 12, 13).

The first result of this analysis is that the human methylome has a large theoretical information capacity. If each CpG site is treated as an independent binary variable, the approximately 30 million CpG sites in the human genome define an upper-bound capacity of approximately 30 Mbit per cell. This value should not be interpreted as the amount of biologically usable methylation information, but rather as the maximal binary state space available under idealized assumptions. When methylation bias is incorporated, using a conservative entropy value of approximately 0.47 bits per CpG, the genome-wide estimate decreases to approximately 14.07 Mbit. Restricting the calculation to the physiological methylation burden equivalent to approximately 18–27 million methylated CpG sites yields an entropy-corrected estimate of approximately 8.44–12.66 Mbit per cell. Thus, even after accounting for biased methylation probabilities, DNA methylation remains a megabit-scale regulatory substrate.

A major implication of this estimate is that DNA methylation should not be viewed merely as a collection of local gene-regulatory switches. Instead, it constitutes a genome-wide regulatory layer with substantial potential information capacity. This does not imply that every methylated cytosine carries independent biological meaning. Rather, it indicates that the methylome provides a large biochemical state space upon which developmental, cellular, environmental, and pathological constraints can act. In this sense, DNA methylation occupies an intermediate position between genome sequence and transient gene expression: it is more stable than most transcriptional fluctuations, but more plastic than DNA sequence itself.

The second major conclusion is that the biologically effective information content of methylation is substantially compressed by methylation bias, spatial correlation, and genomic organization. Neighboring CpGs often behave as coordinated methylation units rather than independent binary variables. When this local correlation is incorporated, the estimated information content decreases substantially. In the present framework, correlation lengths of 5, 10, and 20 CpGs per effective methylation unit reduce the estimate from 14.07 Mbit to approximately 2.81 Mbit, 1.41 Mbit, and 0.70 Mbit, respectively. This range, approximately 0.70–2.81 Mbit per cell, is likely more biologically plausible than the independent-site estimate because it reflects the block-like structure of methylation domains, regulatory elements, CpG islands, shores, enhancers, and partially methylated regions.

This compression is not a weakness of methylation as an epigenetic memory system (7, 17, 18). On the contrary, it is a defining feature of biological information processing. Biological systems rarely store regulatory information as independent, uniformly distributed bits. Instead, information is structured, redundant, error-tolerant, context-dependent, and interpretable through molecular machinery. The methylome is therefore better understood as a compressed regulatory code than as a simple binary storage device. Many CpGs are constitutively methylated or unmethylated, whereas a smaller subset of regulatory regions carries disproportionate information about lineage, developmental state, transcriptional competence, imprinting, or cellular identity.

The stratification of methylation entropy by genomic regulatory class further supports this interpretation. Promoter CpG islands, CpG island shores, enhancers, imprinting control regions, gene bodies, repeats, and transposable elements do not contribute equally to methylation information (2, 6, 11). Promoter CpG islands may often show low variability under normal physiological conditions, but methylation changes at selected promoters can have strong functional consequences. Enhancers and CpG island shores, by contrast, often show greater cell-type specificity and may contribute strongly to identity-defining regulatory programs. Imprinting control regions represent a special case in which a relatively small number of loci can encode highly consequential allele-specific information. Therefore, the biological value of methylation information depends not only on the number of CpGs involved, but also on their genomic location and regulatory interpretation.

The discriminative formulation introduced here provides an additional level of biological constraint. Rather than asking how many methylation states are theoretically possible, mutual information between methylation patterns and cellular identity asks how much methylation reduces uncertainty about the state of the cell. This is a more biologically precise question because it connects methylation information to classification, lineage, and cellular phenotype. In a flat classification problem involving 100 equally probable cell types, the upper bound is only log_2_(100), or approximately 6.64 bits. This value may appear small when compared with megabit-scale genome-wide estimates, but it describes a different quantity: the information required to identify one label among a defined set of alternatives, not the total methylation information encoded across the genome.

The apparent contrast between megabit-scale methylome capacity and bit-scale cell-type discrimination highlights the importance of defining the biological question being asked. Theoretical methylation capacity, entropy-corrected capacity, correlation-corrected capacity, and cell-type-discriminative information are not competing estimates of the same object. They represent different layers of a hierarchy. The theoretical estimate describes the maximal binary state space. The entropy-corrected estimate accounts for methylation bias. The correlation-corrected estimate accounts for local dependency among CpGs. The discriminative estimate asks how much methylation contributes to identifying a biological state. This hierarchy helps reconcile large genome-wide methylation capacity with compact, highly reliable identity codes.

This framework also has implications for disease biology. Aberrant DNA methylation is widely associated with cancer, aging, developmental disorders, metabolic dysfunction, immune dysregulation, and altered cell identity (14–16, 25). However, disease-associated methylation changes are often described qualitatively, as hypermethylated or hypomethylated regions, without a quantitative measure of how much regulatory information has been gained, lost, redistributed, or corrupted. An information-theoretic framework could help distinguish global methylation noise from disease-relevant regulatory reconfiguration. For example, tumors may not only gain or lose methylation at specific loci; they may also alter the entropy, redundancy, correlation structure, or cell-type-discriminative information of the methylome. Quantifying these changes could provide a more rigorous basis for comparing normal and pathological epigenetic states.

The framework proposed here may also be useful for interpreting methylation atlases and single-cell methylome datasets (7, 9, 10). Bulk methylation atlases provide robust cell-type-specific profiles, but they average across cell populations and may obscure rare states or intra-lineage heterogeneity. Single-cell methylation data, although sparse and technically challenging, could allow direct estimation of methylation entropy, mutual information, and regulatory-state diversity at cellular resolution. In future work, the theoretical quantities described here could be refined using empirical methylation distributions, experimentally defined DMRs, chromatin annotations, enhancer maps, transcription factor occupancy, and matched gene expression profiles.

Several limitations should be emphasized. First, the estimates presented here are order-of-magnitude approximations rather than fixed biological constants. The number of CpGs, the fraction of methylated CpGs, the average methylation probability, and the local correlation length vary across tissues, cell types, developmental stages, organisms, and assay platforms. Second, the binary methylated/unmethylated model simplifies a more complex biochemical reality that includes hydroxymethylcytosine and other oxidized derivatives, allele-specific methylation, strand asymmetry, non-CpG methylation in some contexts, and cell-to-cell heterogeneity. Third, the functional interpretation of methylation depends on chromatin state, transcription factor binding, replication timing, three-dimensional genome organization, and gene-regulatory networks. Therefore, methylation information should not be treated as an autonomous code independent of the broader epigenomic system.

Despite these limitations, the present analysis provides a useful conceptual and quantitative foundation. It shows that DNA methylation can be described as an information-bearing molecular system with a large theoretical capacity, a compressed effective regulatory capacity, and a highly constrained cell-type-discriminative component. This layered interpretation avoids two opposing oversimplifications: treating every CpG as an independent functional bit, or dismissing methylation information as small because only a few bits may be required to classify a limited number of cell types. Instead, DNA methylation should be understood as a structured, redundant, and biologically interpreted memory system.

In conclusion, DNA methylation represents a quantifiable epigenetic memory layer whose information content depends on the scale of analysis. At the genome-wide level, the methylome has megabit-scale theoretical and entropy-corrected capacity. At the regulatory level, this capacity is compressed by local CpG correlation and genomic organization. At the phenotypic level, methylation provides discriminative information about cell identity, lineage, and regulatory state. By placing DNA methylation within a Shannon information framework, this study provides a quantitative basis for future analyses of methylation dynamics, epigenetic memory, cell identity, and disease-associated methylation dysregulation. All numerical results shown here are fully reproducible using the Python Modules provided in Supplementary Information S4, which regenerate tables 1, 3 and 4, all information capacity values, and all estimates.

## Supplementary Data statement

Supplementary Data are available at *NAR* Online.

## Data and code availability

All original code has been deposited at https://github.com/JoseCarrascoPujante/Quantifying-the-Information-Capacity-of-DNA-Methylation-as-an-Epigenetic-Memory-System and is publicly available at https://doi.org/10.5281/zenodo.20633508.

## Acknowledgments

The author I.M. was supported by Basque Government funding, grant GIC21/04, and by the Spanish Government, grants PID2024-156800OB-I00 and PID2024-156173OA-I00. JMC acknowledges financial support from Ikerbasque: The Basque Foundation for Science, and from Spanish Ministry of Science (PID2023-148012OB), Spanish Ministry of Health (PI22/01118), and Basque Department of Health (2023111002 & 2022111031).

## Author contributions

IMDF: conceived, designed and directed the investigation, writing – original draft. MF, IM, LM, JC-P, BC-P, LL, GPY, JMC and JIL: investigation, research mapping, writing – review and editing. JC-P Python Modules. All authors participated in writing the final version of the manuscript and agreed to its submission.

