## Supplementary material for "Quantifying the Information Capacity of DNA Methylation as an Epigenetic Memory System": README.md

# Reproducibility package — Information capacity of DNA methylation

Companion code for the manuscript **"Quantifying the Information Capacity of DNA Methylation as an Epigenetic Memory System."** A single Jupyter notebook ([`reproducibility_package.ipynb`](reproducibility_package.ipynb)) regenerates every numerical result and tables 1, 3 (= Table S3.1) and 4 in the article. All output is shown inline within the notebook, no file(s) are generated by it.

[![](https://img.shields.io/badge/License-MIT-yellow.svg)](https://opensource.org/licenses/MIT) [![](https://img.shields.io/badge/License-CC--BY%204.0-lightgrey.svg)](https://creativecommons.org/licenses/by/4.0/) [![DOI](https://zenodo.org/badge/DOI/10.5281/zenodo.20633508.svg)](https://doi.org/10.5281/zenodo.20633508)

## Cite

See CITATION.cff

## Requirements

Python ≥ 3.10 with `numpy` ≥ 1.20, `pandas` ≥ 1.4 and `jinja2` ≥ 3.1.2 (tested with Python 3.13.14, numpy 2.4.2, pandas 3.0.1, jinja2 3.1.6).

## Run

The notebook is intended to run top to bottom. Runtime is a few seconds and no external data or network access is required.

## Reproducibility workflow overview

The notebook reproduces all numerical results in the manuscripts using closed-form Shannon-entropy and Mutual Information expressions. Values are computed in full precision and displayed to two decimals; tables render below each module and can be pasted directly and seamlessly into a word processor. The notebook's reproducibility modules follow the manuscript's reading order so that it can be read and executed side by side with it. Each module asserts its computed results against the manuscript's values, so any change that altered a number would fail loudly.

## Notebook index

| Reproducibility Module | Manuscript item |
|---|---|
| Module 1 | Table 1 — theoretical & entropy-corrected information capacity of DNA methylation (Section 2.1) |
| Module 2 | Table 3 — correlation-corrected effective methylation information (Section 2.4) |
| Module 3 | Table 4 — discriminative methylation information (Section 2.5; equal to Supplementary Information S3 / Table S3.1) |

The full description of each Reproducibility Module is available in **Supplementary Information S4** of the manuscript.

## License

See LICENSE file.
