## Supplementary material for "Quantifying the Information Capacity of DNA Methylation as an Epigenetic Memory System": reproducibility_package.ipynb

```
# %% [markdown]
### Reproducibility Package for the manuscript *Quantifying the Information Capacity of DNA Methylation as an Epigenetic Memory System*.

This standalone notebook regenerates every numerical value and numeric table in
the manuscript from closed-form Shannon-entropy expressions. It needs no
external data or network access, runs top to bottom in a few seconds, and writes
no files: all results, including Word-pasteable tables, are rendered inline.

The published parameters live in a single setup cell. Editing them adapts the
whole framework; the per-module value checks against the manuscript are then
skipped automatically (see `RUN_REGRESSION_CHECKS`), so the notebook still runs.
Full per-cell detailed description is available in Supplementary Information S4 
of the manuscript.

| Module | Manuscript target |
|--------|-------------------|
| 1 | Theoretical, entropy-corrected and burden-adjusted capacity (Table 1) |
| 2 | Correlation-corrected effective methylation units (Table 3) |
| 3 | Discriminative information by cellular-state resolution (Table 4 = Table S3.1) |


**Cite:** Zenodo Concept DOI `10.5281/zenodo.20633508` (resolves to latest version).

# %% [code]
## Environment check and analysis setup - define parameters and functions for 
# information-theoretic calculations and table display.

# Setup and imports
import sys, re, importlib, numpy as np, pandas as pd
pd.set_option('display.float_format', lambda v: f'{v:.2f}')

# ============================================================
#  ENVIRONMENT CHECK
# ============================================================
#  Python
# ============================================================

_py = sys.version_info
assert _py >= (3, 10), f"Python ≥ 3.10 required, got {_py.major}.{_py.minor}"

# ============================================================
# Third-party requirements
# ============================================================

_REQUIRED = {
    "pandas":   (1, 4),      # data manipulation
    "numpy":    (1, 20),     # numerical operations
    "jinja2":   (3, 1, 2),   # HTML report generation
}

def _parse(ver_str):
    "Numeric version, ignoring pre-release suffixes (e.g. '1.3.0rc1')."
    return tuple(int(x) for x in re.findall(r'\d+', ver_str.split('+')[0])[:3])

_ok, _fail = [], []
for _pkg, _min in _REQUIRED.items():
    try:
        _ver_str = importlib.import_module(_pkg).__version__
        _ver = _parse(_ver_str)
        _min_str = ".".join(map(str, _min))
        if _ver >= _min:
            _ok.append(f"  \u2713  {_pkg:<10} {_ver_str}")
        else:
            _fail.append(f"  \u2717  {_pkg:<10} {_ver_str}  (need \u2265 {_min_str})")
    except ImportError:
        _fail.append(f"  \u2717  {_pkg:<10} NOT INSTALLED  (need \u2265 {'.'.join(map(str, _min))})")

# ============================================================
# Report
# ============================================================

print(f"  \u2713  {'Python':<10} {_py.major}.{_py.minor}.{_py.micro}")
print(*_ok, sep="\n")
print()
if _fail:
    print("  FAILED:")
    print(*_fail, sep="\n")
    raise EnvironmentError("Environment check failed; install missing/outdated packages before running.")
else:
    print("All requirements satisfied. Ready to run.")

# The following parameters can be adjusted to accommodate future empirical methylation datasets (Methods 2.11)
N_CPG, P_BIAS = 30e6, 0.9
N_BURDEN = (18e6, 27e6)
LC_VALUES  = [1, 5, 10, 20]
K_VALUES   = [100, 1000, 10000, 100000]

# ============================================================
# Regression-check gate
# ============================================================
# The per-module assertions pin the published manuscript values, acting as a
# reproduction guarantee (Supplementary S4). They must not fire when the
# framework is re-parameterised (Methods 2.11), so we record the published
# defaults and run the checks only when the active parameters are unchanged.
_PUBLISHED_DEFAULTS = {
    "N_CPG": 30e6, "P_BIAS": 0.9, "N_BURDEN": (18e6, 27e6),
    "LC_VALUES": [1, 5, 10, 20], "K_VALUES": [100, 1000, 10000, 100000],
}
RUN_REGRESSION_CHECKS = (
    {"N_CPG": N_CPG, "P_BIAS": P_BIAS, "N_BURDEN": N_BURDEN,
     "LC_VALUES": LC_VALUES, "K_VALUES": K_VALUES} == _PUBLISHED_DEFAULTS
)
if not RUN_REGRESSION_CHECKS:
    print("Parameters differ from published defaults; regression checks skipped.")


# ============================================================
# Define core functions and 2-decimal table display helper
# ============================================================
def shannon_entropy(p):
    p = np.asarray(p, float)
    with np.errstate(divide='ignore', invalid='ignore'):
        h = -p*np.log2(p) - (1-p)*np.log2(1-p)
    return np.where((p == 0) | (p == 1), 0.0, h)

def capacity_mbit(n, h):   return n * float(h) / 1e6    # information capacity, Mbit/cell
def effective_units(n, lc): return n / lc / 1e6         # independent methylation units, millions
def discriminative_bits(k): return float(np.log2(k))    # log2(K), bits; k = number of distinguishable methylation-defined cellular states

def show_table(df, caption):
    floats = df.select_dtypes('float').columns
    return df.style.format('{:.2f}', subset=floats).hide(axis='index').set_caption(caption)

# %% [code]
## Reproducibility Module 1 - theoretical and entropy-corrected DNA methylation capacity (Section 2.1 & Table 1)
h05, h09 = float(shannon_entropy(0.5)), float(shannon_entropy(P_BIAS))
lo, hi = capacity_mbit(N_BURDEN[0], h09), capacity_mbit(N_BURDEN[1], h09)
t1 = pd.DataFrame({
    'Model': ['Theoretical maximum, p = 0.5', 'Biased methylation, p = 0.9',
             'Physiological methylated CpG range, 18-27 million CpGs'],
    'Entropy per CpG site (bits)': [h05, h09, h09],
    'Estimated capacity per cell (Mbit)': [f'~{capacity_mbit(N_CPG, h05):.2f}',
                                 f'~{capacity_mbit(N_CPG, h09):.2f}', f'~{lo:.2f}-{hi:.2f}']})

# Assertions to verify that the calculated values match manuscript values (rounded to 2 decimal places)
if RUN_REGRESSION_CHECKS:
    assert round(capacity_mbit(N_CPG, h05), 2) == 30.00
    assert round(capacity_mbit(N_CPG, h09), 2) == 14.07
    assert (round(lo, 2), round(hi, 2)) == (8.44, 12.66)

# Display the final table for theoretical maximum and entropy-corrected DNA methylation information capacity
show_table(t1, 'Table 1. Information capacity of DNA methylation')

# %% [code]
## Reproducibility Module 2 - correlation-corrected effective methylation units (Section 2.4 & Table 3)
t3 = pd.DataFrame({
    'Average local correlation length L<sub>c</sub> (CpGs per unit)': LC_VALUES,
    'Effective independent units, N<sub>eff</sub> (millions)': [effective_units(N_CPG, lc) for lc in LC_VALUES],
    'Correlation-corrected information (Mbit)': [capacity_mbit(N_CPG/lc, h09) for lc in LC_VALUES]})

# Assertions to verify that the calculated values match manuscript values (rounded to 2 decimal places)
if RUN_REGRESSION_CHECKS:
    assert [round(v, 2) for v in t3.iloc[:, 1]] == [30.00, 6.00, 3.00, 1.50]
    assert [round(v, 2) for v in t3.iloc[:, 2]] == [14.07, 2.81, 1.41, 0.70]

# Display the final table for correlation-corrected methylation information
show_table(t3, 'Table 3. Correlation-corrected methylation information')

# %% [code]
## Reproducibility Module 3 - discriminative methylation information (Section 2.5 & Table 4 = Table S3.1)
t4 = pd.DataFrame({
    'Biological classification space': ['Broad cell-type atlas', 'Fine cellular subtypes',
        'Cell type × tissue context', 'Cell type × tissue × developmental or functional state'],
    'Number of distinguishable states K': K_VALUES,
    'Upper-bound discriminative information (bits)': [discriminative_bits(k) for k in K_VALUES]})

# Assertion to verify that the calculated values match manuscript values (rounded to 2 decimal places)
if RUN_REGRESSION_CHECKS:
    assert [round(v, 2) for v in t4.iloc[:, 2]] == [6.64, 9.97, 13.29, 16.61]

# Display the final table for discriminative information
show_table(t4, 'Table 4. Upper-bound discriminative information as a function of cellular-state resolution')
```
