## Supplementary material for "Quantifying the Information Capacity of DNA Methylation as an Epigenetic Memory System": Supplementary Information.pdf

DNA methylation is one of the most extensively studied forms of epigenetic regulation. It consists of the enzymatic addition of a methyl group to DNA bases, most commonly to the fifth carbon of cytosine, generating 5-methylcytosine. In animals, this modification occurs predominantly at CpG dinucleotides, whereas in plants methylation is also found in CHG and CHH sequence contexts, where H represents A, T, or C. Importantly, DNA methylation does not alter the underlying nucleotide sequence. Instead, it provides an additional regulatory layer through which cells can modulate chromatin organization, transcriptional activity, genome stability, and cellular identity (1, 2).

The relevance of DNA methylation as an epigenetic memory system derives from its capacity to persist through cell division while remaining potentially reversible. Once established, methylation patterns can be copied after DNA replication, allowing daughter cells to inherit regulatory information from the parental cell. This feature distinguishes DNA methylation from transient transcriptional responses and allows it to participate in the long-term stabilization of developmental programs. In this sense, methylation marks function not merely as biochemical modifications, but as molecular records of cellular history, lineage commitment, and environmental exposure.

The establishment and propagation of DNA methylation require coordinated enzymatic activities. De novo methyltransferases generate new methylation patterns at previously unmethylated sites, particularly during development, differentiation, and genome defense. Maintenance methyltransferases then preserve these patterns after DNA replication by recognizing hemimethylated DNA and restoring methylation on the newly synthesized strand. Through this replication-coupled copying mechanism, a methylation state can be transmitted across many mitotic divisions, thereby contributing to the stable inheritance of gene expression programs.

At the same time, DNA methylation is not an irreversible mark. Demethylation can occur passively when methylation is not restored after replication, leading to progressive dilution across cell divisions. It can also occur actively through enzymatic pathways that remove or chemically modify methylated cytosines. In mammals, active demethylation is closely associated with TET-mediated oxidation of 5-methylcytosine and subsequent DNA repair processes. In plants, active demethylation is mediated by DNA glycosylases that excise methylated cytosines and initiate base excision repair. Thus, methylation memory is best understood as a dynamic equilibrium between deposition, maintenance, and erasure.

A central property of DNA methylation is its cell-type specificity. Although nearly all somatic cells within an organism share the same genome, they exhibit distinct methylation landscapes that reflect their developmental origin, functional specialization, and transcriptional state. Promoters, enhancers, gene bodies, repetitive elements, and transposable elements can all display different methylation profiles depending on the cellular context. These patterns help define which genes remain active, silent, poised, or protected from inappropriate activation. Consequently, methylation contributes to the emergence and maintenance of cell identity without requiring changes in DNA sequence.

The regulatory function of DNA methylation depends strongly on genomic context. At many gene promoters, especially those associated with CpG islands in mammals, high methylation is generally linked to transcriptional repression. By contrast, methylation within gene bodies can correlate with transcriptional activity or with the suppression of cryptic transcription initiation. In repetitive regions and transposable elements, DNA methylation plays a major role in transcriptional silencing, thereby protecting genome integrity. These context-dependent effects show that methylation cannot be interpreted as a simple universal on/off signal; rather, its meaning depends on local sequence features, chromatin state, transcription factor occupancy, developmental stage, and cell type.

DNA methylation is widespread across life, being present in bacteria, archaea, fungi, plants, and animals (3–5). However, its biological roles have diversified across evolution. In prokaryotes, DNA methylation is often involved in host defense, DNA replication control, and restriction-modification systems. In eukaryotes, it has become deeply integrated into chromatin regulation, transposon repression, imprinting, X-chromosome inactivation, developmental control, and cellular differentiation.

The model plant Arabidopsis thaliana has been an especially important test bench for advancing our understanding of DNA methylation. Its genome is relatively compact (∼ 95% smaller than that of mammals), with about ∼10.5% of cytosines methylated (6). The A. thaliana methylome was the first to be fully mapped at single-base resolution in 2006 (7–10).

Taken together, these features justify viewing DNA methylation as a form of epigenetic memory. Its stability allows cells to preserve regulatory states over time, while its reversibility permits developmental reprogramming, adaptation, and response to internal or external perturbations. DNA methylation therefore occupies an intermediate position between genetic information, which is highly stable, and transcriptional activity, which is often rapidly fluctuating. It provides a semi-stable regulatory memory layer that links genome sequence, chromatin architecture, cellular identity, environmental history, and metabolic state.

### Supplementary Information S2. Molecular mechanisms of maintenance and erasure

DNA methylation is not a static chemical imprint but a dynamically maintained regulatory state. Its persistence depends on the balance between three partially overlapping processes: de novo methylation, which establishes new cytosine methylation marks; maintenance methylation, which copies pre-existing methylation patterns after DNA replication; and demethylation, which removes or dilutes methylation marks either actively through enzymatic pathways or passively through replication-dependent loss. Together, these processes allow DNA methylation to behave as a stable yet reversible form of cellular memory.

In mammals, cytosine methylation occurs predominantly at CpG dinucleotides and is catalyzed by DNA methyltransferases. DNMT3A and DNMT3B are generally regarded as the main de novo methyltransferases, establishing methylation patterns during early development, lineage specification, and cellular differentiation. DNMT1, by contrast, is the canonical maintenance methyltransferase. After DNA replication, newly synthesized daughter strands are initially unmethylated, generating hemimethylated CpG sites. DNMT1 recognizes these hemimethylated substrates and restores symmetrical methylation, thereby propagating the methylation pattern through mitotic cell divisions. This copying process is essential for preserving cell identity and transcriptional programs across proliferating somatic lineages (11).

Maintenance methylation in mammals is also strongly dependent on chromatin context. DNMT1 does not act as an isolated enzyme scanning naked DNA; rather, it is recruited and regulated by replication-associated and chromatin-associated factors. UHRF1 plays a central role in this process by recognizing hemimethylated CpGs and histone marks associated with repressive chromatin, thereby helping target DNMT1 to recently replicated DNA. This coupling between DNA methylation, histone modifications, nucleosome organization, and replication timing explains why methylation memory is not simply sequence-intrinsic, but embedded within a broader epigenetic architecture. Recent work further indicates that DNMT3 enzymes can contribute not only to de novo methylation but also to the long-term preservation of methylation landscapes, especially in genomic regions where DNMT1-mediated copying alone is insufficient (12).

The erasure of DNA methylation in mammals occurs through both passive and active mechanisms. Passive demethylation takes place when methylation marks are not restored after DNA replication, leading to progressive dilution of 5-methylcytosine across successive cell divisions. Active demethylation, in contrast, is initiated by the ten-eleven translocation enzymes TET1, TET2, and TET3, which oxidize 5-methylcytosine to 5-hydroxymethylcytosine and further derivatives such as 5-formylcytosine and 5-carboxylcytosine. These oxidized bases can either serve as relatively stable regulatory marks in some contexts or be processed through DNA repair pathways, ultimately restoring unmodified cytosine. Thus, mammalian demethylation is not a single-step reversal but a multistage biochemical pathway that links epigenetic regulation with DNA oxidation and repair (13).

In plants, DNA methylation is mechanistically more diverse because cytosine methylation occurs in three sequence contexts: CG, CHG, and CHH, where H represents A, T, or C. Each context is maintained by partially distinct enzymatic systems. CG methylation is primarily maintained by MET1, the plant functional counterpart of mammalian DNMT1. CHG methylation is maintained mainly by chromomethylases such as CMT3, in close interaction with histone H3 lysine 9 methylation. CHH methylation, which is asymmetric and therefore cannot be copied directly after replication, requires continuous de novo activity mediated by DRM2 through the RNA-directed DNA methylation pathway, as well as CMT2 activity in heterochromatic regions. These mechanisms are especially important for transposon silencing and genome stability (14).

The RNA-directed DNA methylation pathway provides plants with a sequence-specific mechanism for establishing and reinforcing methylation. In this pathway, small interfering RNAs guide DRM2-containing methylation complexes to homologous genomic regions, allowing methylation to be targeted by RNA-DNA sequence complementarity. This system is particularly relevant for transposable elements, repetitive DNA, and heterochromatic regions, but it can also influence nearby genes and regulatory elements. Because RdDM can continuously re-establish methylation at selected loci, plant methylation memory is maintained not only by copying existing marks but also by RNA-guided surveillance of the genome (15).

Plant demethylation is also more directly linked to base excision repair than in mammals. Active DNA demethylation is mediated by the REPRESSOR OF SILENCING 1 family of bifunctional DNA glycosylases, including ROS1, DEMETER, DML2, and DML3. These enzymes recognize and excise 5-methylcytosine from DNA, creating an abasic site that is subsequently repaired to restore an unmethylated cytosine. This pathway allows plants to actively counterbalance methyltransferase activity and prevent excessive methylation spreading into regulatory regions. The coexistence of methylation and demethylation activities creates a dynamic equilibrium in which methylation boundaries can be maintained with locus-specific precision (14).

In addition, in plants, methylation is maintained by several enzyme families. These include the MET1 methyltransferases (16), chromomethylases (17), and DRM1/2 domains-rearranged methyltransferases (18). Demethylation is driven by glycosylases such as DEMETER (19) and DEMETER-like enzymes DML2 and DML3 (20). The RNA-directed DNA methylation pathway (RdDM) further establishes sequence-specific marks (21), stabilized by global “methylstat” feedback (22), and by interactions between methyltransferase classes (23). In mammals, transcription factors central to pluripotency, including OCT4, SOX2, and NANOG, participate in guiding methylation states (24).

This balance between maintenance and erasure is central to the interpretation of DNA methylation as epigenetic memory. A methylation pattern can persist only if maintenance mechanisms compensate for dilution during replication and resist inappropriate demethylation. Conversely, developmental reprogramming, environmental responses, aging, and disease can shift the balance toward methylation gain or loss. Therefore, DNA methylation memory should not be viewed as a permanently fixed molecular record, but as a metastable regulatory state continuously reconstructed by enzymatic activity, chromatin context, transcriptional signals, DNA repair, and cellular metabolism.

In this sense, the methylome resembles a regulated dynamic memory system. Its stability derives from feedback between DNA methyltransferases, demethylases, histone modifications, small RNAs, transcription factors, and replication-coupled maintenance pathways. Its plasticity derives from the ability of the same system to erase, remodel, or overwrite methylation patterns during development and in response to internal or external perturbations. This dual property, high stability under normal lineage propagation and controlled reversibility during reprogramming, explains why DNA methylation can function both as a mechanism of cellular identity and as a substrate for epigenetic adaptation.

#### Supplementary Information S3: Hierarchical classification of methylation-defined cellular states and sensitivity analysis of discriminative information

##### S3.1. Discriminative information with 100 equally probable cell types

In the main text, discriminative methylation information was formulated as the mutual information between methylation patterns and cellular identity. This formulation follows the Shannon information-theoretic interpretation of information as reduction of uncertainty (25). Let M denote the methylation representation and C denote the cellular identity variable:

$$I(M;C) = H(C) - H(C|M)$$

Because mutual information is bounded by the entropy of the variable being decoded,

$$I(M;C) <= H(C)$$

the numerical magnitude of the discriminative information depends strongly on how cellular identity is defined. This point is particularly important because modern methylation atlases and single-cell epigenomic studies show that methylation patterns can discriminate cellular identity at multiple biological resolutions, from broad cell classes to fine regulatory states (1, 26, 27).

If C is represented as a flat classification variable containing 100 equally probable cell types, the maximal discriminative information is:

$$H(C) = log_2(100) \approx 6.64 \text{ bits}$$

This value should not be interpreted as the total amount of information stored in the methylome. Rather, it is the upper bound for a coarse classification task: identifying one cell-type label among 100 predefined alternatives. A more biologically realistic representation of cellular identity may require a hierarchical or combinatorial definition, in which a cell is described not only by broad cell type, but also by tissue origin, developmental lineage, subtype, activation state, differentiation state, disease-associated state, or allele-specific regulatory status. Such hierarchical representations are consistent with current cell taxonomy and ontology frameworks, which emphasize that cell identity includes both cell types and cell states (28, 29).

Thus, the cellular identity variable can be generalized as:

$$C^* = (C_{type}, C_{tissue}, C_{lineage}, C_{subtype}, C_{state}, C_{imprinting})$$

where C* represents a methylation-defined regulatory cellular state rather than a single flat cell-type label.

#### S3.2. Hierarchical expansion of the cellular identity space

The entropy of a hierarchical cellular identity variable can be decomposed as:

$$H(C^*) = H(C_{type}) + H(C_{tissue}|C_{type}) + H(C_{lineage} \mid C_{tissue}) + H(C_{subtype}|C_{lineage}) + H(C_{state} \mid C_{subtype})$$

This expression emphasizes that the discriminative information available to methylation depends on the biological resolution at which cellular identity is decoded. A broad cell-type atlas may define approximately 100 distinguishable categories, whereas a higher-resolution atlas incorporating tissue origin, cellular subtypes, activation states, developmental stages, and disease-associated regulatory states may define many more methylation-distinguishable cellular states. Large-scale cell ontology resources and human cell atlas initiatives explicitly motivate this hierarchical view of cellular identity (28, 29).

For K equally probable methylation-defined cellular states, the maximal discriminative information is:

$$H(C) = log_2(K)$$

where K is the number of distinguishable cellular or regulatory states. This simple sensitivity analysis does not assume that all such states have already been experimentally resolved by methylation alone. Instead, it shows how the upper bound of I(M;C) changes when the decoded biological state space is represented at different resolutions.

#### S3.3. Sensitivity analysis of discriminative information

The following Table S3.1 shows how the upper bound of discriminative methylation information increases as the number of distinguishable methylation-defined cellular states increases.

**Table S3.1. Sensitivity analysis of discriminative methylation information as a function of cellular-state resolution.**

| Biological classification space | Number of distinguishable states K | Upper-bound discriminative information $H(C)= log_2$ (K) |
|---|---|---|
| Broad cell-type atlas | 100 | 6.64 bits |
| Fine cellular subtypes | 1,000 | 9.97 bits |
| Cell type × tissue context | 10,000 | 13.29 bits |
| Cell type × tissue × developmental or functional state | 100,000 | 16.61 bits |

This sensitivity analysis shows that the value of 6.64 bits is not an intrinsic upper bound of DNA methylation as an information-bearing system. It is the upper bound imposed by a low-resolution classification problem involving 100 equally probable labels. When cellular identity is defined at higher biological resolution, the entropy of the decoded state space increases accordingly.

The hierarchical formulation provides a bridge between genome-wide methylation information and cell-type-discriminative methylation information. Genome-wide methylation entropy can reach the megabit scale because it reflects the number of possible methylation states distributed across millions of CpG sites or effective methylation units. In contrast, discriminative information between cellular states is bounded by the number of biological categories being decoded.

Therefore, the relatively small value obtained for 100 cell types should not be interpreted as evidence that DNA methylation carries little biological information. Instead, it indicates that only a small number of reliable bits is required to identify one class among a limited set of alternatives. In biological systems, such compact discriminative information can be highly meaningful if it robustly distinguishes cell identity, developmental lineage, regulatory state, or disease-associated phenotype.

This interpretation is consistent with the view of DNA methylation as a compressed regulatory code. Most CpG sites do not act as independent functional bits. Instead, methylation information is structured into regulatory domains, differentially methylated regions, enhancer-associated CpGs, CpG island shores, imprinting control regions, and lineage-defining methylation markers. Large epigenomic reference maps and functional genomics projects support the view that regulatory information is strongly dependent on genomic context, chromatin state, and cell type (30–32).

Several classes of regulatory elements may be especially relevant to methylation-defined state spaces. Enhancers can retain epigenetic memory of developmental history, whereas CpG island shores have been shown to display tissue-specific methylation changes and disease-associated alterations (33, 34). These observations support the idea that methylation-defined cellular states may include tissue origin, developmental memory, regulatory context, and pathological remodeling, rather than only broad cell-type labels.

##### S3.4. Relationship with methylation-defined regulatory states

A methylation-defined cellular state may be broader or narrower than a conventional cell type. For example, two cells classified under the same broad cell type may differ in developmental maturity, activation status, tissue microenvironment, disease state, or epigenetic age. Conversely, different cell types may share methylation features inherited from a common lineage. For this reason, the discriminative variable C should be interpreted flexibly.

In a coarse model:

$$C = Ctype$$

where Ctype represents broad cell identity.
In a higher-resolution model:

$$C = C^*$$

where C* represents a combinatorial cellular regulatory state.
Under this framework, the discriminative information estimate becomes:

$$I(M;C^*) <= H(C^*)$$

rather than being restricted to:

$$I(M;Ctype) <= H(Ctype)$$

This distinction is important because methylation patterns may encode regulatory information that exceeds broad cell-type classification, including tissue-specific enhancer states, lineage memory, imprinting status, developmental transitions, and disease-associated epigenetic remodeling. It is also relevant for epigenome-wide association and biomarker studies, where cell-type heterogeneity can dominate methylation variability and must be considered when interpreting disease-associated signals (35).

In summary, the discriminative information carried by DNA methylation depends on the resolution of the biological identity variable being decoded. For a flat classification of 100 cell types, the upper bound is approximately 6.64 bits. However, if methylation-defined cellular identity is represented as a hierarchical or combinatorial state space, the upper bound increases to approximately 9.97 bits for 1,000 states, 13.29 bits for 10,000 states, and 16.61 bits for 100,000 states. Thus, the 6.64-bit estimate should be interpreted as a lower-resolution discriminative benchmark, not as the total information capacity of DNA methylation. DNA methylation can simultaneously display megabit-scale genome-wide entropy and compact bit-scale discriminative information, depending on whether the analysis asks how many methylation states are possible or how many biological states need to be distinguished.

To clarify the interpretation of the numerical estimates used throughout the manuscript, Supplementary Table S3.2 summarizes the assumptions, quantities measured, and main limitations of each layer of the proposed framework

#### Supplementary Table S3.2. Model assumptions and interpretation of methylation information estimates

| Estimate | Main assumption | What it measures | Main limitation |
|---|---|---|---|
| **Theoretical maximum** | CpG sites are treated as independent binary variables with maximal entropy (p = 0.5). | Upper-bound binary methylation state space of the human methylome. | Biologically unrealistic because CpGs are biased, locally correlated, and context-dependent. |
| **Entropy-corrected capacity** | CpG methylation is represented as a biased binary variable with an assumed average methylation probability, for example p = 0.9. | Statistical entropy of methylation states after accounting for methylation bias. | Does not account for spatial correlation, regulatory genomic context, or functional interpretation. |
| **Methylation-burden-adjusted approximation** | The physiological methylation burden is approximated as a range equivalent to approximately 18–27 million methylated CpG sites. | A burden-adjusted estimate of methylation-state capacity within a physiologically plausible methylation range. | This is not a direct count of independent informative CpG variables and should not be interpreted as biologically decoded information. |
| **Correlation-corrected capacity** | Neighboring CpGs are grouped into effective methylation units according to an assumed average local correlation length, Lc. | Effective number of statistically independent methylation units after local CpG coupling. | The correlation length is an order-of-magnitude parameter and may vary across tissues, cell types, genomic regions, and assay platforms. |
| **Regulatory-class stratified entropy** | CpGs or effective methylation units are grouped by genomic regulatory class, such as CpG islands, shores, enhancers, gene bodies, repeats, and imprinting control regions. | Context-dependent methylation entropy across regulatory genomic compartments. | Requires empirical annotation and class-specific methylation distributions; functional consequences are not uniform across loci. |
| **Discriminative information** | Cellular identity is treated as a defined biological-state variable, and methylation | Mutual information or upper-bound uncertainty reduction about cell type, lineage, | Depends strongly on how cellular states are defined and does not represent the |

| | | | |
|---|---|---|---|
| | patterns reduce uncertainty about that variable. | tissue context, developmental state, or disease-associated state. | total information capacity of the methylome. |

This table summarizes the assumptions, interpretation, and main limitations of each information-theoretic estimate used in the methylation-capacity framework. The estimates should be interpreted as order-of-magnitude quantities under explicitly defined assumptions, rather than fixed biological constants.

### Supplementary Information S4: Reproducible Python modules for methylation information estimates

#### Overview

The Jupyter notebook regenerates every numerical result reported in the main text from closed-form information-theoretic expressions. It requires no external data or network access, runs top to bottom in a few seconds, and follows the manuscript's reading order. All quantities are computed in full double precision and rounded only for display (two decimals). When the notebook runs with the published default parameters, each module asserts its computed values against the manuscript, so any change that altered a published result halts execution; when the parameters are modified for adaptation (Methods 2.11), these regression checks are skipped automatically via the RUN_REGRESSION_CHECKS flag, leaving the computational modules fully functional. The notebook consists of one header cell, one setup cell, and three reproducibility modules. The three reproducibility modules calculate the numerical values in Tables 1, 3 and 4 (= Table S3.1) of the manuscript using Python Standard Library modules and the packages Numpy, Pandas and Jinja.

#### Environment check, parameters, and core functions.

This cell first validates the runtime environment, confirming Python ≥ 3.10 and the presence of numpy (≥ 1.20), pandas (≥ 1.4) and jinja2 (≥3.1.2) versions, and aborting with an explicit message if any requirement is unmet, to prevent downstream errors. It then defines, as a single source of truth, the parameters used throughout: the reference number of CpG sites ($N_{CpG} = 30 \times 10^6$), the conservative per-CpG methylation probability (p = 0.9), the physiological methylated-CpG range ($18 – 27 \times 10^6$), the set of local correlation lengths ($L_c$ = 1, 5, 10, 20), and the set of cellular-state resolutions (K = $10^2$, $10^3$, $10^4$, $10^5$). Finally, it defines the core functions: the binary Shannon entropy, $H(p) = -p \cdot log_2(p) - (1-p) \cdot log_2(1-p)$, which calculates the bits per CpG, with H = 0 at p = 0 or 1; the information capacity $I = N \cdot H$, reported in megabits per cell; the effective number of independent units $N_{eff} = N_{CpG}/L_c$, and the discriminative bound $log_2(K)$. Last, a small helper function is defined that renders each result table as a formatted, caption-bearing HTML table suitable for direct transfer to a word processor. The setup cell also records the published defaults and sets RUN_REGRESSION_CHECKS, a boolean that is true only when the active parameters equal those defaults; each module's value assertions are guarded by this flag.

#### Reproducibility Module 1. Upper-bound methylome capacity and Physiological methylated CpG range (Section 2.1, Table 1).

Computes the theoretical and entropy-corrected information capacity of the methylated DNA. At maximal entropy (p = 0.5, H = 1.00 bit) the $30 \times 10^6$ CpG sites give a theoretical maximum of ~30.00 Mbit per cell; under the biased model (p = 0.9, H = ~0.47 bit) the entropy-corrected estimate is ~14.07 Mbit per cell; restricting to the physiological methylated range ($18 – 27 \times 10^6$ CpGs) yields ~8.44 to 12.66 Mbit per cell. The cell assembles these values into Table 1.

#### Reproducibility Module 2. Correlation-corrected effective units (Section 2.4, Table 3).

Applies the correlation correction $I_{corr} = \left(N_{CpG}/L_c\right) \cdot H(p)$, which accounts for neighbouring CpGs behaving as coordinated methylation blocks. For $L_c$ = 1, 5, 10, 20 the effective number of independent units ($N_{eff}$) is 30.0, 6.0, 3.0, and 1.5 million, giving correlation-corrected information of 14.07, 2.81, 1.41, and 0.70 Mbit per cell, respectively. The cell assembles these into Table 3.

#### Reproducibility Module 3. Discriminative information between cell states (Section 2.5, Table 4; equal to Supplementary Table S3.1).

Computes the upper bound on the discriminative information that methylation can carry about cellular identity, $I(M; C) \leq H(C) = log_2(K)$. For K = $10^2$, $10^3$, $10^4$, $10^5$ number of equally probable distinguishable cellular states this gives 6.64, 9.97, 13.29, and 16.61 bits, respectively. The cell assembles these values into Table 4.
